## Supplementary material for "Hierarchical semantic composition of biosimulation models using bond graphs": S1 Table

| | Parameter | Value/Option | Description |
|---|---|---|---|
| Function coefficients | $a_1$ | 0.0005422 | |
| | $b_1$ | 0.1317 | |
| | $c_1$ | 0.03315 | |
| | $a_2$ | 0.0003591 | |
| | $b_2$ | 0.181 | |
| | $c_2$ | 0.06554 | |
| Fit options | $DiffMinChange$ | $1.0e^{-3}$ | |
| | $DiffMaxChange$ | 0.1 | |
| | $Robust$ | $LAR$ | Minimizing the Least Absolute Residuals. |
| | $Algorithm$ | $Levenberg - Marquardt$ | If the fit generated by the trust-region algorithm is not acceptable, try Levenberg-Marquardt. |
| | $MaxFunEvals$ | 60,000 | Maximum number of allowed evaluations for the selected function. |
| | $TolFun$ | $1.0e^{-9}$ | Termination tolerance in stopping conditions for the function. |
| | $TolX$ | $1.0e^{-10}$ | Termination tolerance in stopping conditions for the coefficients. |
