## Supplementary figures and images for "Hierarchical semantic composition of biosimulation models using bond graphs"

### S2 Figure

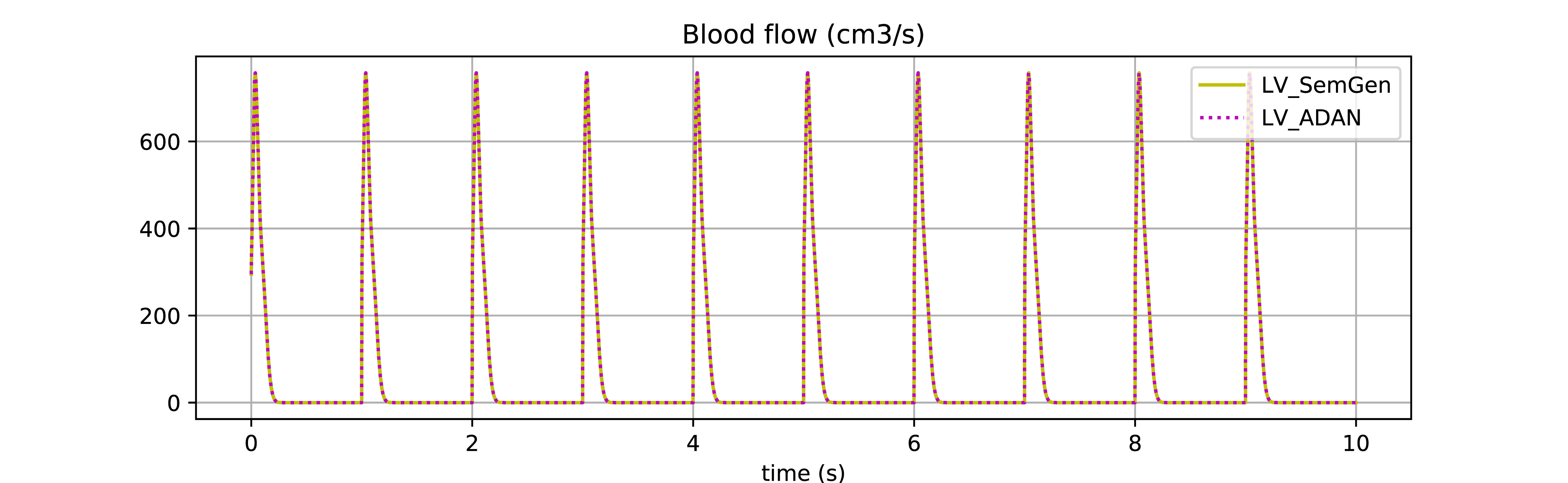
